# Replication-competent HIV-1 in human alveolar macrophages and monocytes despite nucleotide pools with elevated dUTP

**DOI:** 10.1101/2022.05.03.490432

**Authors:** Junru Cui, Mesfin Meshesha, Natela Churgulia, Christian Merlo, Edward Fuchs, Jennifer Breakey, Joyce Jones, James T. Stivers

## Abstract

Although CD4^+^ memory T cells are considered the primary latent reservoir for HIV-1, replication competent HIV has been detected in tissue macrophages in both animal and human studies. During *in vitro* HIV infection, the depleted nucleotide pool and high dUTP levels in monocyte derived macrophages (MDM) leads to proviruses with high levels of dUMP, which has been implicated in viral restriction or reduced transcription depending on the uracil base excision repair (UBER) competence of the macrophage. Incorporated dUMP has also been detected in viral DNA from circulating monocytes (MC) and alveolar macrophages (AM) of HIV infected patients on antiretroviral therapy (ART), establishing the biological relevance of this phenotype but not the replicative capacity of dUMP-containing proviruses. As compared to *in vitro* differentiated MDM, AM from normal donors had 6-fold lower levels of dTTP and a 6-fold increased dUTP/dTTP, indicating a highly restrictive dNTP pool for reverse transcription. Expression of uracil DNA glycosylase (UNG) was 8-fold lower in AM compared to the already low levels in MDM. Accordingly, ∼80% of HIV proviruses contained dUMP, which persisted for at least 14-days due to low UNG excision activity. Unlike MDM, AM expression levels of UNG and SAM and HD domain containing deoxynucleoside triphosphate triphosphohydrolase 1 (SAMHD1) increased over 14 days post-HIV infection, while dUTP nucleotidohydrolase expression decreased. These AM-specific effects suggest a restriction response centered on excising uracil from viral DNA copies and increasing relative dUTP levels. Despite the restrictive nucleotide pools, we detected rare replication competent HIV in AM, peripheral MC, and CD4^+^ T cells from ART-treated donors. These findings indicate that the potential integration block of incorporated dUMP is not realized during *in vivo* infection of AM and MC due to the near absence of UBER activity. In addition, the increased expression of UNG and SAMHD1 in AM post-infection is too slow to prevent integration. Accordingly, dUMP persists in integrated viruses, which based on *in vitro* studies, can lead to transcriptional silencing. This possible silencing outcome of persistent dUMP could promote viral latency until the repressive effects of viral dUMP are reversed.

## INTRODUCTION

Although CD4^+^ T cells are the primary targets of HIV, myeloid cells such as monocytes and macrophages are infected by R5 tropic and dual tropic strains of HIV and have unique properties that could lead to persistent HIV infection even in the presence of active retroviral therapy (ART)^1, 2^. For instance, macrophages have distinct and malleable metabolic properties compared to T cells that make them intrinsically resistance to HIV-induced cytopathic effects and less susceptible to some antiretroviral drugs^3, 4^. These properties include, but are not limited to, innate immunity pathways that involve viral nucleic acid lethal mutation by enzymatic cytidine deamination^5, 6^, dramatically reduced dNTP pools and composition through the action of SAMHD1 dNTPase^7, 8^, and the ability to transiently access G0 and G1 stages of the cell cycle^4, 9, 10^. In addition, recent reappraisals of macrophage biology indicate that these cells can have multi-year life spans in certain tissue environments and possess self-renewal properties that are similar to those of memory T cells^11–15^. These considerations have led to the still debated conclusion that myeloid lineage cells serve as a reservoir for HIV, even in the presence of ART^16^.

The experimental evidence supporting the proposal that macrophages serve as a persistent HIV reservoir in humans after initiation of ART is extensive, but less than definitive. In part, the continuing uncertainty arises from the relative difficulty in obtaining tissue macrophages in sufficient numbers to detect and quantify rare viral nucleic acid, excluding the possibility that contamination by infected or phagocytized T cells accounts for any positive result, and the use of different viral detection assays by various researchers, each with different limits-of-detection or specificity (see the complete review by Wong *et al*)^12, 13^. Nevertheless, reports from different research groups over the past twenty years have consistently (but not uniformly) detected viral RNA, proviral DNA and p24 antigen in isolated monocytes and tissue macrophages from HIV infected patients on ART^17–21^. In many cases, convincing controls for T cell contamination were performed and genetic analysis of viral progeny indicated distinct genotypes for HIV produced from macrophages as compared to T cells from the same patient^18, 22^. In addition, studies using animal models such as SIV-infected macaques^23, 24^, humanized BLT mice and myeloid-only mice^25, 26^ have provided support for the contention that HIV persistence in the presence of ART can be partially attributed to infected myeloid cells, especially in the CNS where ART is less effective^27, 28^.

Three phenotypic traits of *in vitro* differentiated monocyte-derived macrophages (MDM) that are distinct from CD4^+^ T cells are their overall low dNTP pool levels^4, 29^, elevated [dUTP]/[TTP]^20, 30–32^, and resting cell (non-dividing) status^30^. The depleted nucleotide pool promotes a potentially restrictive environment for HIV infection^29^, but the resting cell phenotype allows MDM to persist after infection by avoiding the apoptotic responses observed with infected T cells^28^. The perturbed nucleotide pool leads to delayed cDNA synthesis for *in vitro* differentiated MDM and incorporation of large amounts of dUMP in viral DNA products during reverse transcription^20, 32^. Further, because MDM have low expression levels of uracil base excision repair (UBER) enzymes^20, 30, 31^, dUMP can persist in proviral DNA and has been associated with transcriptional repression of viral genes20,31,34,35.

This simple picture of non-dividing MDM being repair-deficient has been complicated by recent findings that these cells exist in two populations: (i) a major G0 population where the dNTP pool is depleted, the DNA repair activity is low and HIV proviruses are heavily uracilated^20^, and (ii) a minor pseudo-G1 population characterized by a normal dNTP pool, DNA repair activity, and low levels of repair-associated DNA replication^4, 9, 36^. The pseudo-G1 state can be transiently and reversibly accessed from the G0-state, providing a mechanism for long-lived macrophages to repair genomic DNA, but also an increased chance of viral infection while in the pseudo-G1 state. The presence of two populations with greatly different susceptibilities to infection complicates quantitative evaluation of the effects of depleted and imbalanced dNTP pools^9, 20^. The presence of G0 and pseudo-G1 populations is but one example of the malleable nature of the macrophage phenotype that is determined by the microenvironment of the tissue in which the cell resides^36^. Importantly, the nucleotide pool and DNA repair attributes of *in vitro* differentiated MDM have never been compared to *in vivo* differentiated tissue macrophages. Nevertheless, abundant viral dU/A pairs have been detected in HIV infected monocytes and alveolar macrophages isolated from patients receiving ART suggesting that similar nucleotide pool compositions exist for *in vitro* and *in vivo* differentiated macrophages^20^.

Here we report the first evaluation of the dNTP pool status of *in vivo* differentiated alveolar macrophages (AM) obtained from normal and HIV-infected donors using ART. Our findings confirm that the phenotype of depleted dNTP pools and elevated dUTP/TTP extends to *in vivo* differentiated AM and that the expression of the uracil base excision enzyme uracil DNA glycosylase (UNG) is vanishingly low in these cells, which promotes the persistence of dUMP in proviral DNA. Since dU/A base pairs have previously been associated with suppression of RNAPII transcription^20, 31, 34^, uracilated proviral DNA may be a unique transcriptional silencing mechanism in macrophages. We also report using a quantitative viral outgrowth assay (QVOA) that replication competent HIV virus can be isolated from *in vivo* infected AM and peripheral blood monocytes.

## RESULTS

### Alveolar macrophages (AM) have very low dNTP pools, high dUTP/TTP and depleted uracil DNA glycosylase (UNG)

Using a highly sensitive ddPCR assay we previously reported the detection of HIV DNA in peripheral MC from 6 out of 6 fully ART-suppressed HIV patients (50-100 pol copies/10^6^ cells)^20^. Further, 5/6 of the patient MC were positive for dUMP content, as were two AM samples obtained from a single patient pre- and post-ART^20^. Importantly, the viral uracilation phenotype was unique to viral DNA obtained from MC and AM, but was absent in CD4 T cells^20, 31^. These results established that (i) HIV DNA was detectable in myeloid cells using the most sensitive detection method, and (ii) viral DNA from these cells was uniquely marked by the presence of dUMP. However, these studies did not assess whether the viral DNA was integrated or whether replication competent viruses were present.

To further characterize the dNTP pool and UBER phenotypes of AM, we purified AM obtained by bronchoalveolar lavage from non-infected donors (**Table 1**). The AM were purified as described in Methods, and the absence of contaminating T cells was established using RT-qPCR targeting the TCRβ mRNA, which is selectively expressed in T cells^39^. From this analysis, we confirmed that T cells were present at <1 T cell/10^2^ AM. Further, ninety-five percent of the isolated AM were CD68 positive and 90% were functional as judged by a positive phagocytosis assay employing pHrodo *E. coli* cells (**Fig. 1a, b**).

**Figure 1.**
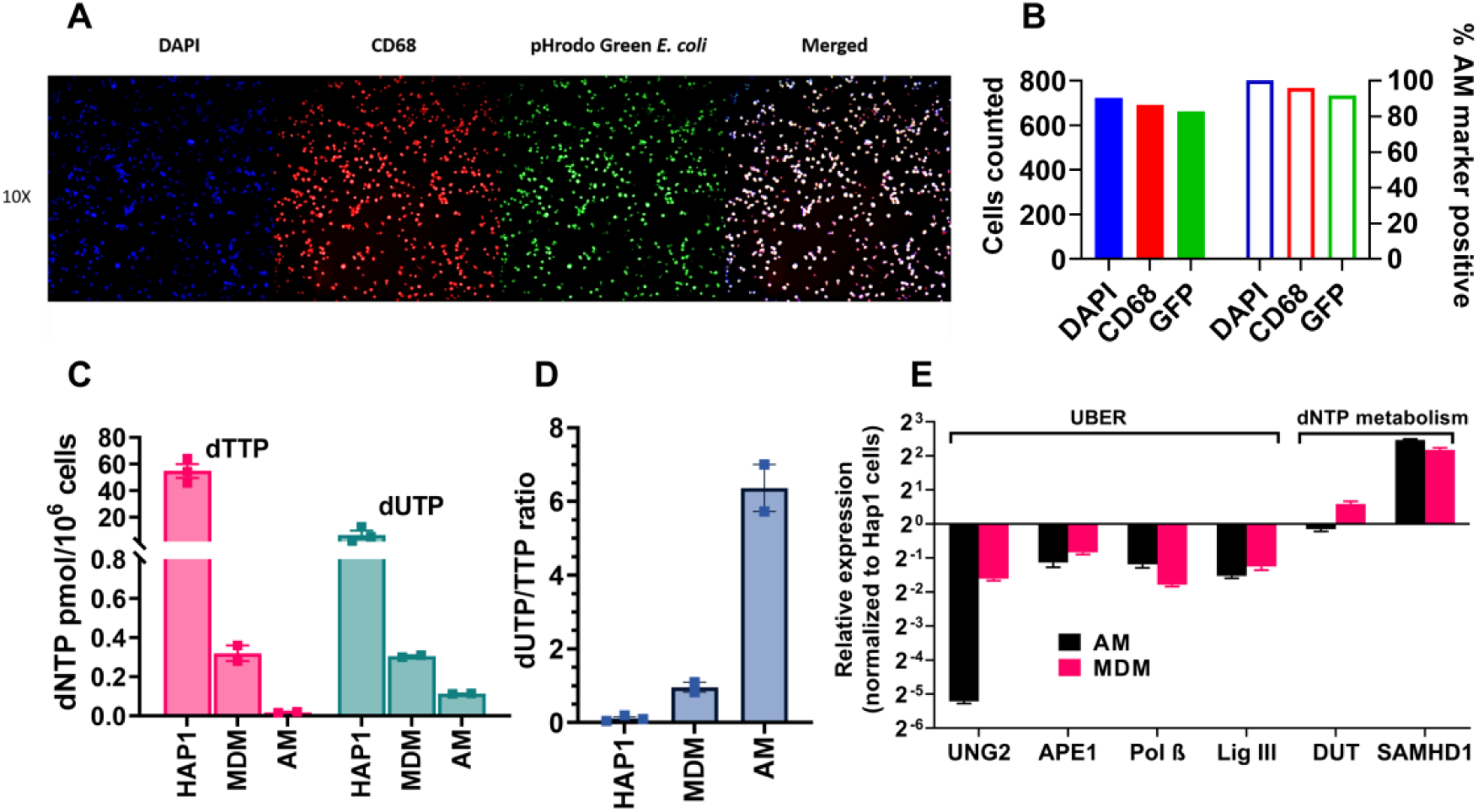
Confirmation of in vivo AM phenotype and comparison of dTTP, dUTP nucleotide and UBER enzyme levels with that of MDM. (**A**) Fluorescence microscopy of AM stained with DAPI DNA stain (blue), CD68 cell surface marker (red) and pHrodo Green *E. coli* metabolic marker (green). An enlarged merged image is shown on the right. (**B**) AM cells in panel A were counted using ImageJ and the percent of DAPI stained cells that were also stained with the other three markers was calculated. (**C**) dUTP and dTTP levels in extracts from dividing HAP1 cells, non-dividing MDM, and AM cells as determined using a single nucleotide extension assay (SNE)(Fig. S1). The *in vitro* differentiated MDM and the *in vivo* differentiated AM cells were from the same healthy donors. (**D**) dUTP/dTTP ratio in extracts from dividing HAP1 cells, non-dividing MDM, and AM cells. (**E**) Baseline mRNA expression levels of proteins involved in uracil base excision repair (UBER) and dNTP metabolism in AM and MDM cells. Expression levels were normalized to dividing HAP1 cells for comparison purposes. 18S ribosomal RNA was used as the calibration standard in all measurements. Error bar represents SD, the mean was calculated with three biological replicates for HAP1 cells, and two health donors for AM and MDM data.

**Table 1.**
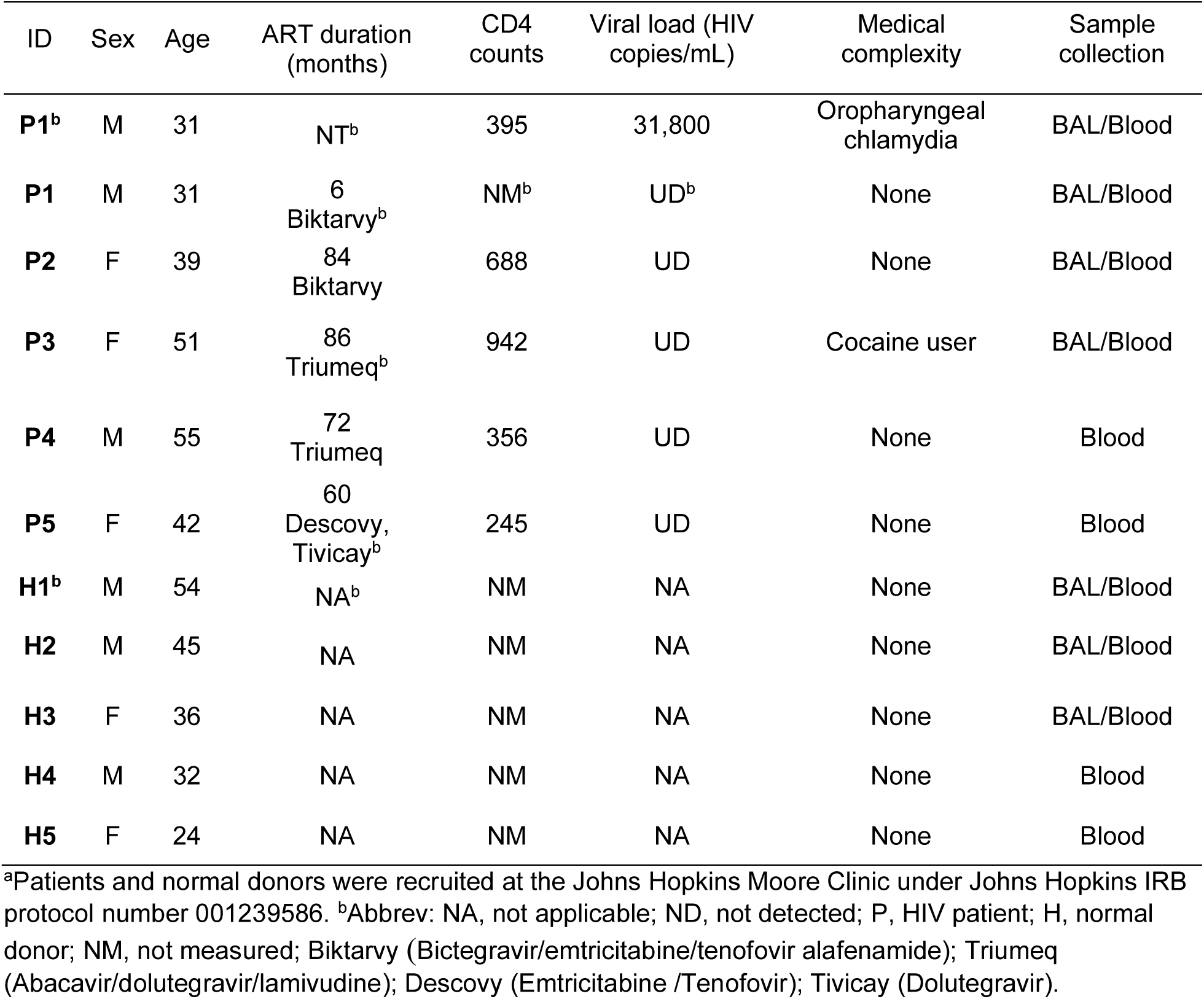
Characteristics of study participants^a^.

| ID | Sex | Age | ART duration (months) | CD4 counts | Viral load (HIV copies/mL) | Medical complexity | Sample collection |
| --- | --- | --- | --- | --- | --- | --- | --- |
| **P1<sup>b</sup>** | M | 31 | NT<sup>b</sup> | 395 | 31,800 | Oropharyngeal chlamydia | BAL/Blood |
| **P1** | M | 31 | 6<br>Biktarvy<sup>b</sup> | NM<sup>b</sup> | UD<sup>b</sup> | None | BAL/Blood |
| **P2** | F | 39 | 84<br>Biktarvy | 688 | UD | None | BAL/Blood |
| **P3** | F | 51 | 86<br>Triumeq<sup>b</sup> | 942 | UD | Cocaine user | BAL/Blood |
| **P4** | M | 55 | 72<br>Triumeq | 356 | UD | None | Blood |
| **P5** | F | 42 | 60<br>Descovy,<br>Tivicay<sup>b</sup> | 245 | UD | None | Blood |
| **H1<sup>b</sup>** | M | 54 | NA<sup>b</sup> | NM | NA | None | BAL/Blood |
| **H2** | M | 45 | NA | NM | NA | None | BAL/Blood |
| **H3** | F | 36 | NA | NM | NA | None | BAL/Blood |
| **H4** | M | 32 | NA | NM | NA | None | Blood |
| **H5** | F | 24 | NA | NM | NA | None | Blood |

<sup>a</sup>Patients and normal donors were recruited at the Johns Hopkins Moore Clinic under Johns Hopkins IRB protocol number 001239586. <sup>b</sup>Abbrev: NA, not applicable; ND, not detected; P, HIV patient; H, normal donor; NM, not measured; Biktarvy (Bictegravir/emtricitabine/tenofovir alafenamide); Triumeq (Abacavir/dolutegravir/lamivudine); Descovy (Emtricitabine /Tenofovir); Tivicay (Dolutegravir).

We used a modified single-nucleotide primer extension assay to assess the dUTP and dTTP levels in AM for comparison with *in vitro* differentiated MDM and dividing HAP1 human cells (**Fig. 1c and Supplemental Fig. S1**)^30, 31^. As compared to the HAP1 cell line, the dTTP levels were reduced by 2,700 and 170-fold in AM and MDM, respectively. Accordingly, the dUTP/dTTP ratio in AM is 32-fold greater than the HAP1 line, and more than 6-fold greater than the already elevated ratio seen in MDM (**Fig. 1d**). These measurements of high dUTP levels in AM aligns with our previous Ex-ddPCR measurements of high uracil content in HIV DNA isolated from infected AM (**Fig. 1c**)^20^.

We then used RT-qPCR to probe the base line expression levels of key UBER enzymes (UNG, APE1, pol β, lig III, DUT) and the dNTPase SAMHD1 in AM and MDM prior to infection with HIV (**Fig. 1e**). For comparative purposes, we normalized the measured expression levels to those found in the HAP1 dividing cell line. These measurements established that UNG, the first enzyme in the UBER pathway that excises uracil bases from DNA, was present at an 8-fold lower level in AM as compared to MDM and about 50-fold less than the HAP1 line. The other UBER enzymes showed similar expression levels in AM and MDM, which were about 2 to 4-fold less than HAP1 cells. As expected, both AM and MDM showed a 4-fold greater expression of SAMHD1 compared to the HAP1 cells. Taken together, these data show that AM have extraordinarily low dNTP levels, elevated dUTP/dTTP and greatly reduced UNG activity. Thus, dUMP incorporated into viral DNA during infection of AM would be expected to persist at higher levels than MDM due to the severe depletion of UNG and dTTP.

### HIV infection kinetics of AM and MDM and relative dUMP levels in viral DNA products

There have been very limited studies of HIV infection of AM or any other primary tissue macrophages. To ascertain the kinetics and relative efficiency of HIV infection of AM and to examine how many viral DNA products contain detectable dUMP, we infected AM and MDM with an equivalent number of R5 tropic HIV^Bal^ virus (MOI = 1) and monitored viral DNA products and viral p24 levels over 21 days (**Fig. 2**). Using PCR primers designed to detect early (ERT) and late (LRT) reverse transcription DNA products we found that the ERT levels peaked at 3 copies per cell at 1-day post infection of AM while the LRT copies reached a final level of about 0.5 copies per cell (**Fig. 2a**). Thus, only about one-sixth of the ERT produced in AM were converted to more mature viral DNA products, suggesting a restrictive step early in infection of AM. In contrast, the ERT and LRT levels in MDM increased steadily over the course of infection, peaking at 4 copies per cell at 7-days post infection. By day 14, both the ERT and LRT copies in MDM diminished by almost a factor of two, suggesting a late restrictive step not observed with AM (**Fig. 2a**).

**Figure 2.**
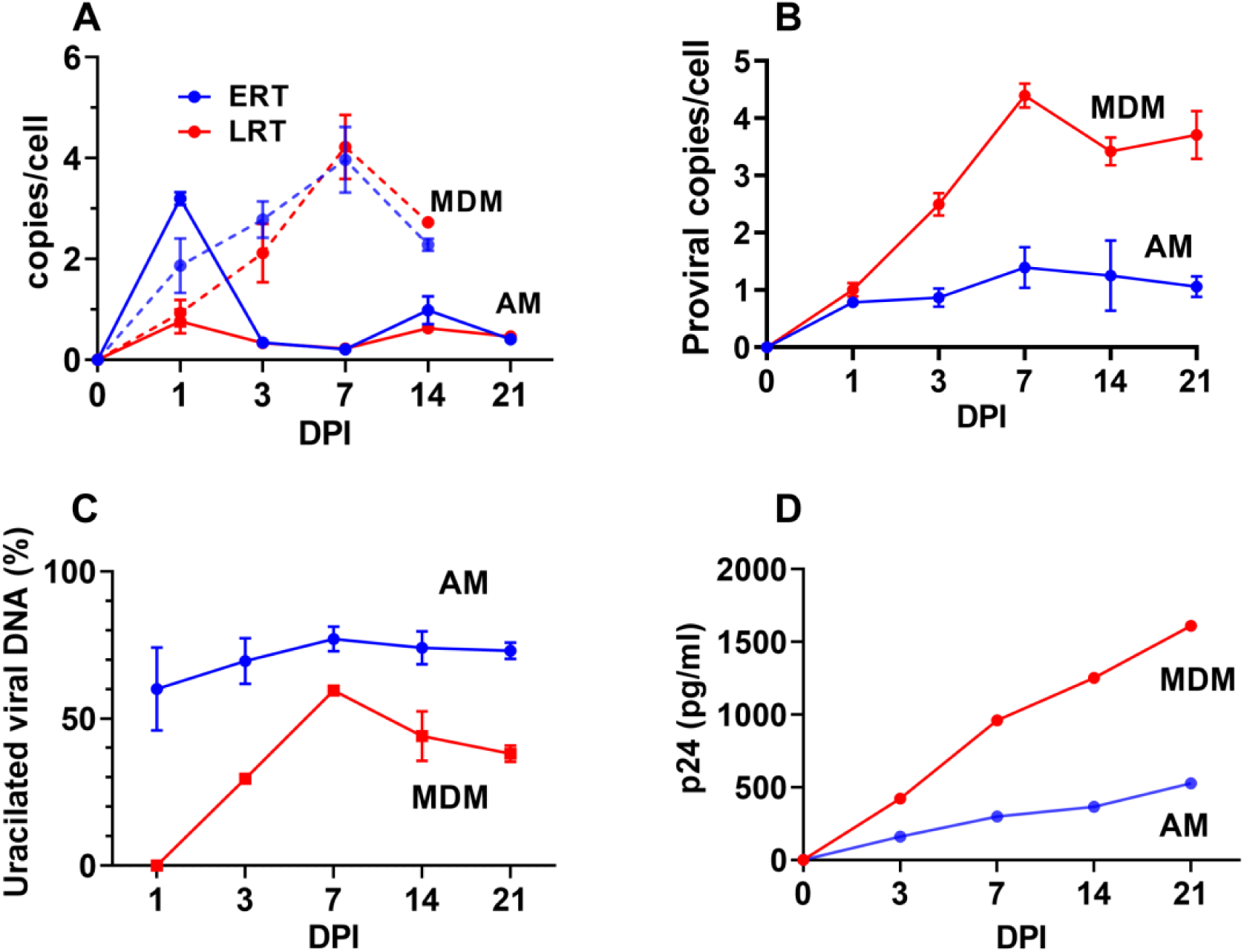
Comparison of *in vitro* infection of AM and MDM with HIV^Bal^. (**A**) Kinetics for appearance of early (ERT) and late (LRT) reverse transcripts in AM and MDM as determined by RT-qPCR. The RPP30 gene was used to calculate the average transcript copies per cell. Sequences and probes are listed in Table S1. (**B**) Kinetics for appearance of proviral DNA copies in MDM and AM as determined by alu-gag qPCR. (**C**) Percentage of proviruses present in MDM and AM that contain dUMP as determined by alu-gag Ex-qPCR. (**D**) Viral protein 24 (p24) levels in culture supernatants of infected AM and MDM were measured by ELISA. The supernatant p24 levels are from identical infections (MOI = 1) using equal numbers of plated target cells (50,000 per well in 96-well plate). Errors are means with standard deviations, N = 2 for A, B, C and N = 1 for D.

Using the Alu-gag qPCR method for amplification of integrated viruses, we found that proviral copies of infected AM reached a plateau of about one copy per cell after about 7 days and remained stable out to day 21 (**Fig. 2b**). In contrast, proviral copies of infected MDM increased steadily from one copy per cell at 1-day post infection to four copies at 7-days and remained roughly the same out to 21-days post infection (**Fig. 2b**). Thus, AM and MDM showed distinct kinetics for viral integration.

We used the uracil-excision qPCR (Ex-qPCR) method to determine the fraction of LTR DNA products that contained dUMP from 1 to 21 days post-infection of AM and MDM (**Fig. 2c**). For AM, viral dUMP was present in ∼80% of the LTR products at day one and the level was unchanged over the course of the infection. In contrast, MDM showed no detectable dUMP in LTR DNA products at day 1 and the levels increased between days 3 and 7 up to 60% of viral LTR copies. Unlike AM, the fraction of LTR copies containing dUMP decreased to about 40% in MDM between days 7 and 21 (**Fig. 2c**). The absence of detectable dUMP in viral DNA at one day post-infection of MDM has been observed consistently and attributed to rapid infection of the MDM cell population present in the pseudo-G1 state where dUTP is not present^40^. The slow reduction in dUMP over time has been attributed to the slow replacement of proviral dUMP with TMP by base excision repair^40^. The early appearance of dUMP and its persistence in AM suggests that the pseudo-G1 population is a minor contribution, and that uracil excision activity is significantly reduced compared to MDM (consistent with the UNG expression profiling).

The p24 levels in culture supernatants of infected AM and MDM steadily increased over 21 days for both MDM and AM (**Fig. 2d**), with the final p24 level at day 21 about three-fold higher for MDM, which is similar in magnitude to the greater proviral copy number for MDM (**Fig. 2b**). Taken together, these data indicate slower infection kinetics, decreased infection efficiency, and increased dUMP persistence for AM as compared to MDM.

### UBER enzyme expression levels in AM and MDM post-HIV infection

We measured large changes in the expression levels of UNG and two other enzymes in our panel at various times after infection of AM with R5 tropic HIV^Bal^ virus (**Fig. 3**). Strikingly, AM showed a 20-fold induction of UNG mRNA between days 3 and 14 post infection, while no significant increases in UNG expression were observed with MDM over the same time frame. In addition, the expression level of SAMHD1 was increased ∼30-fold in AM and dUTPase showed a rapid 16-fold decrease in expression by day 3 (**Fig. 3a**). These changes in SAMHD1 and dUTPase expression were not observed for MDM, which instead showed an ∼30-fold increase in expression of DNA cytidine deaminase A3A, but not A3G (**Fig. 3b**). The post-infection upregulation of UNG and SAMHD1 and the downregulation of dUTPase suggest a coordinated response to the invading virus. Specifically, these changes would be expected to impede viral replication via increased SAMHD1 dNTPase activity and other SAMHD1-dependent mechanisms^37^, increasing the already high dUTP levels by decreasing dUTPase activity, and promoting excision of viral uracils by increasing UNG activity. However, these changes in expression levels in AM may not be rapid enough to impact the early stages of viral infection in a significant way. Further, the increased levels of UNG appear to be ineffective at removing proviral uracils over 21 days post-infection (**Fig. 2c**). This effect may indicate that integrated viruses with uracil are protected by chromatin compaction or other mechanisms.

**Figure 3.**
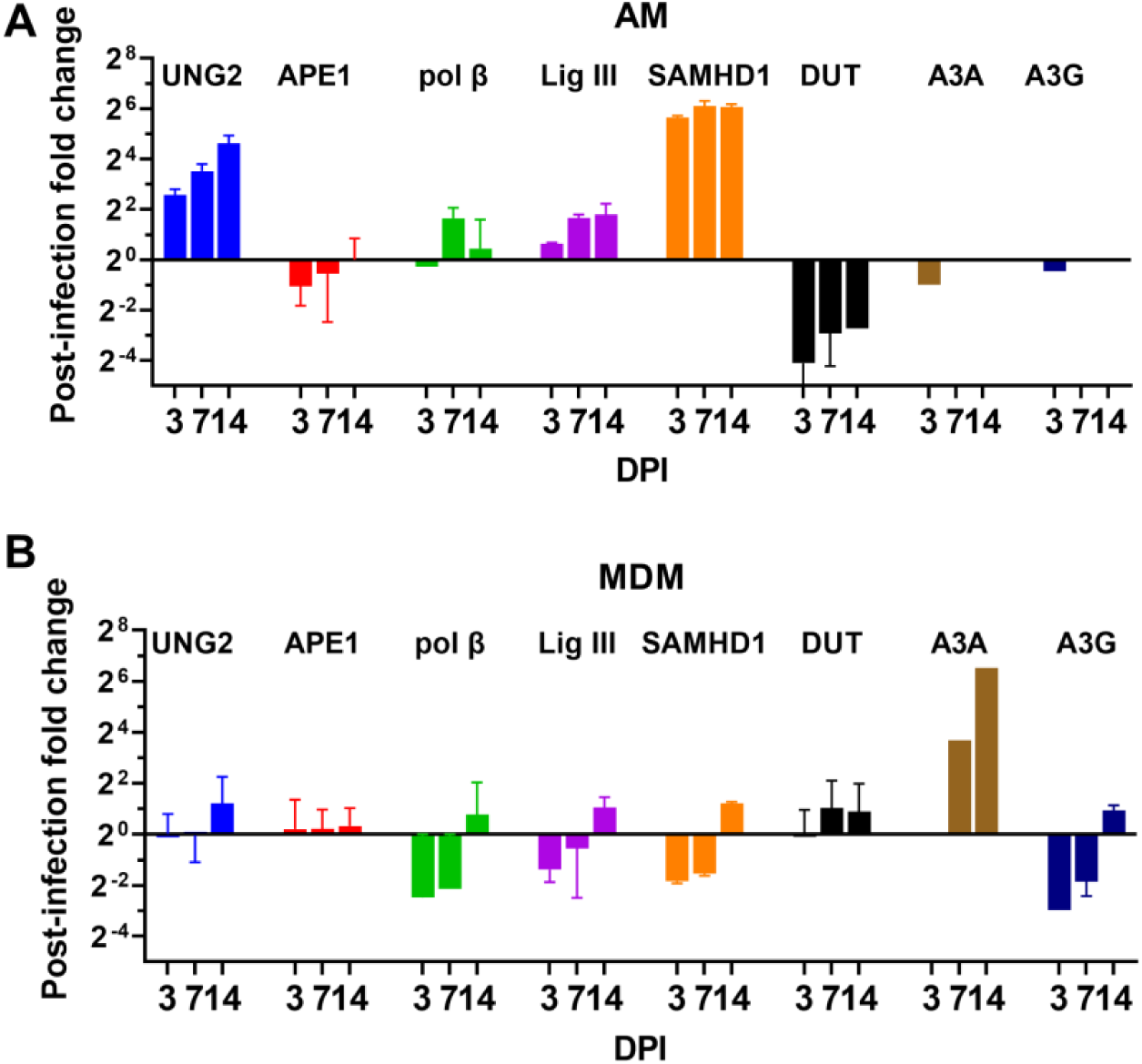
UBER gene expression in AM and MDM post-infection with HIV^Bal^. (**A**) RT-qPCR was used to measure the time-dependent changes in mRNA levels of the indicated enzymes post-infection of AM. (**B**) Time-dependent changes in mRNA levels of the indicated enzymes post-infection of MDM. AM data was collected from one healthy donor and the errors are from three technical replicates. The MDM data is the average from three healthy donors. The fold-change in copy number is relative to the levels before infection and the 18S ribosomal RNA expression was used as the calibration standard for all samples. All primers are listed in Table S1. Errors are means with standard deviations, N = 2 for AM and N = 3 for MDM (where N is number of donors).

### Quantification of replication competent virus present in AM, MC and T cells from ART-suppressed patients

An optimized QVOA assay for macrophages and monocytes was used to determine the number of productively infected myeloid and T cells obtained from five patients (**Table 2**)(**Fig. 4a**)^23^. A total of 2.6 to 15.0 million myeloid cells (AM or MC) and 3.0 to 20.0 million T cells were examined for each donor, which sets the limit of detection for each cell type (**Table 2**). The QVOA experiments with AM and MC were assessed for the presence of TCRβ mRNA using RT-qPCR and the percent contamination of T cells in each QVOA was not greater than 0.9% (AM) or 1.7% (MC) (**Fig. 4b**). Since replication competent viruses are present at a frequency of less than 1 per million CD4^+^ T cells (see below), the probability of infected CD4^+^ T cells harboring replication competent virus in the AM and MC QVOA reactions was 0.02 and 0.04 per million AM and MC, respectively. Accordingly, this level of T cell contamination cannot account for any of the positive AM and MC QVOA reactions (see below).

**Figure 4.**
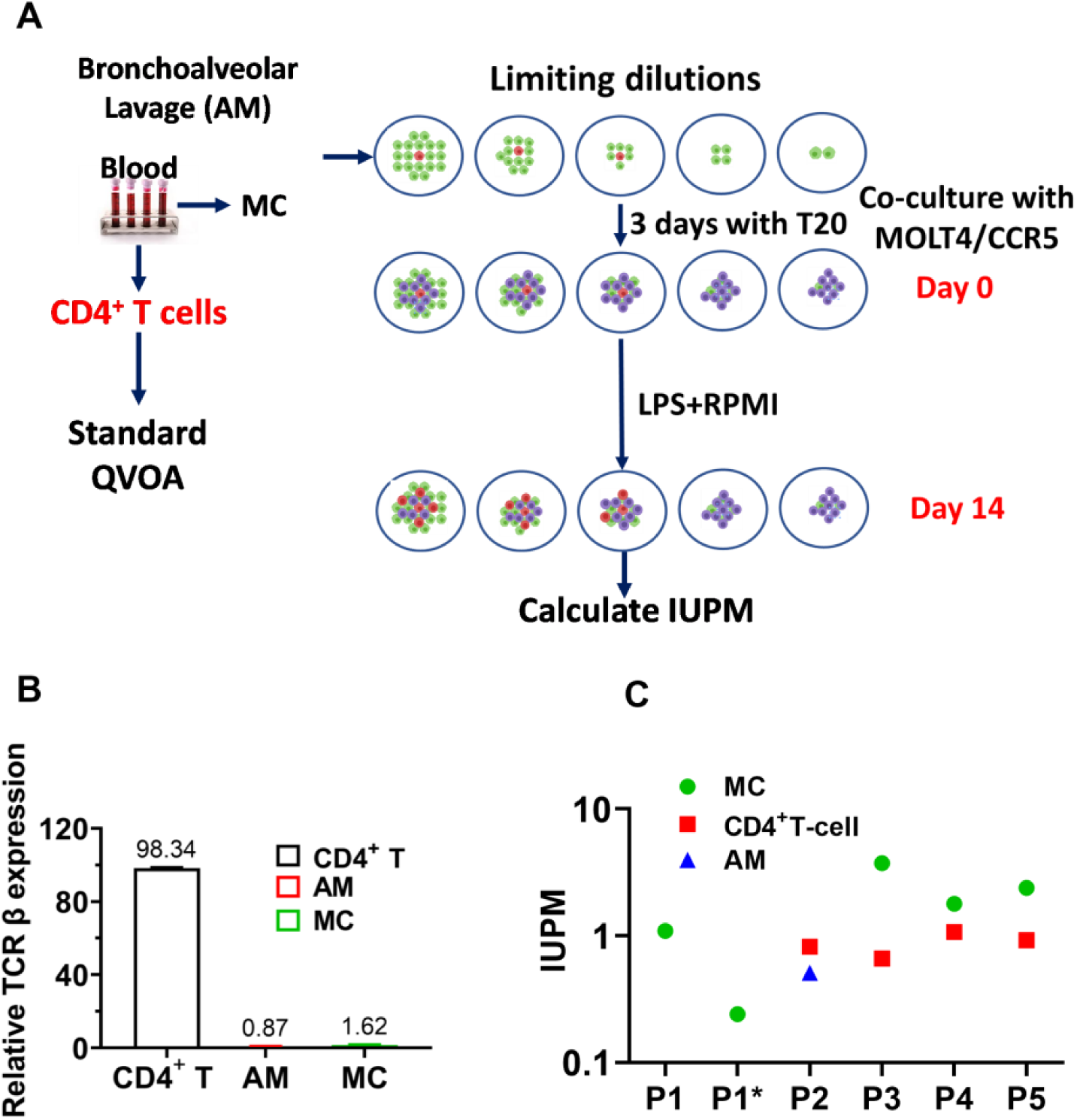
Replication competent HIV-1 detected by QVOA from AM, MC and T cells of HIV patients under suppressive ART. (**A**) Monocytes and CD4^+^ T cells from blood and AMs from BAL fluid were collected from five HIV-infected patients and purified by T cell pan isolation kit. T cells were first cultured for three days using stimulating conditions and then plated in serial dilutions into three to five wells. Monocytes purified by the pan monocyte isolation kit were checked for T-cell contamination by RT-qPCR using TCR-β primer pairs (98.38 ± 0.43 % purity). MDM were generated by culturing MC under adherent conditions for seven days in the presence of M-CSF, and the cells were serially diluted into three to five wells. AM were plated directly in 3 to 5 serial dilution wells. Cells were cultured in the presence of the HIV protease inhibitor T20 (Enfuvirtide) as indicated. Nonadherent cells and T20 were removed prior to activation with LPS and coculturing with MOLT4/CCR5 cells. (**B**) TCR β RNA levels detected in QVOA lysates of T, MC and AM cells were measured by RT-qPCR using a seven-point standard curve established by serial dilution. The relative expression percent was normalized to CD4^+^ T cells. The mean is the average of three donors. Errors are means with standard deviations, N = 3 for AM, T and MC, where N is number of donors. (**C**) The infectious units per million cells (IUPM) of AM, MC and CD4^+^ T cells was determined by RT-qPCR using the Infection Frequency Calculator which utilizes limiting dilution Poisson statistics and the number of positive wells and the input number of cells (https://silicianolab.johnshopkins.edu).

**Table 2.**
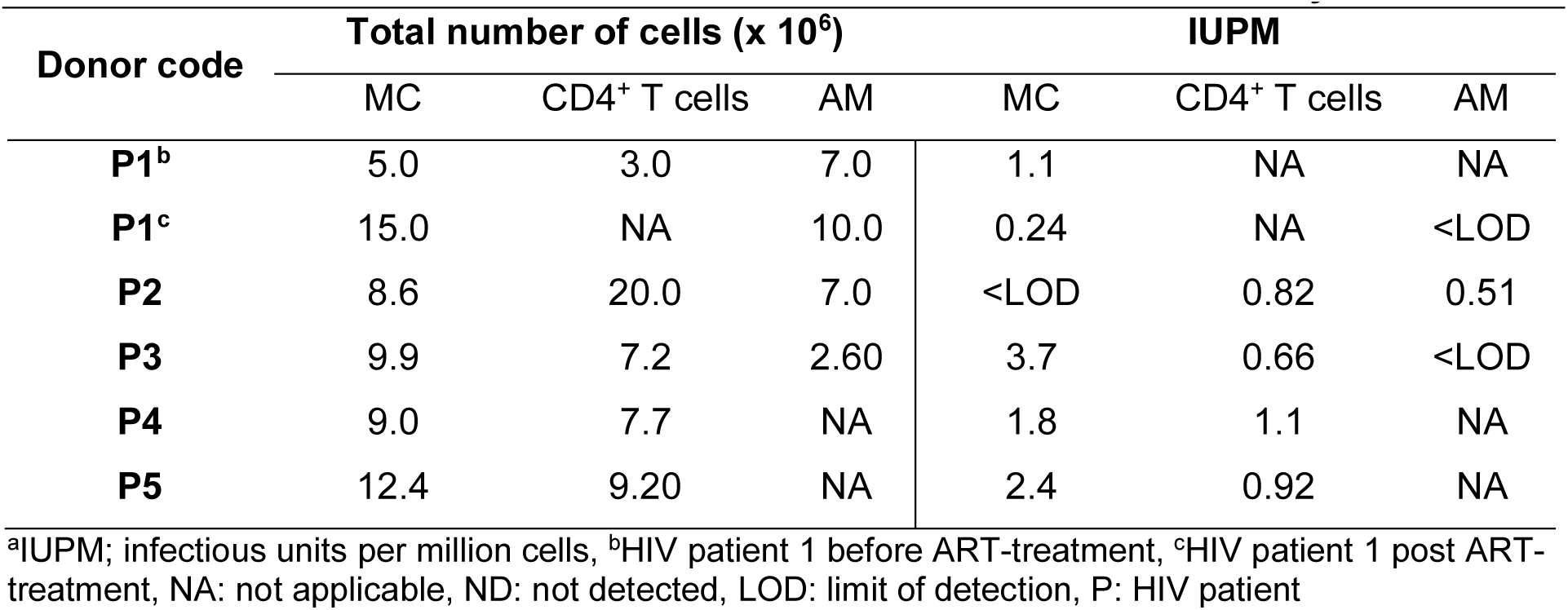
Total number of cells assessed and calculated IUPM^a^ in this study.

| Donor code | Total number of cells (x 10<sup>6</sup>) |  |  | IUPM |  |  |
| --- | --- | --- | --- | --- | --- | --- |
|  | MC | CD4<sup>+</sup> T cells | AM | MC | CD4<sup>+</sup> T cells | AM |
| **P1<sup>b</sup>** | 5.0 | 3.0 | 7.0 | 1.1 | NA | NA |
| **P1<sup>c</sup>** | 15.0 | NA | 10.0 | 0.24 | NA | <LOD |
| **P2** | 8.6 | 20.0 | 7.0 | <LOD | 0.82 | 0.51 |
| **P3** | 9.9 | 7.2 | 2.60 | 3.7 | 0.66 | <LOD |
| **P4** | 9.0 | 7.7 | NA | 1.8 | 1.1 | NA |
| **P5** | 12.4 | 9.20 | NA | 2.4 | 0.92 | NA |

<sup>a</sup>IUPM; infectious units per million cells, <sup>b</sup>HIV patient 1 before ART-treatment, <sup>c</sup>HIV patient 1 post ART-treatment, NA: not applicable, ND: not detected, LOD: limit of detection, P: HIV patient

Using qPCR, we detected viral RNA in the QVOA supernatants originating from one AM sample out of three that were analyzed and from four of the five MC QVOA experiments (**Table 2** and **Fig. 4b**). In addition, viral RNA was detected in four out of five T cell QVOA experiments. In these experiments, QVOA wells that contained at least 50 RNA copies/mL were deemed positive and the limit of detection was 10 HIV RNA copies per reaction. The number of latently infected cells that contained replication-competent virus was then determined using an algorithm for maximum likelihood estimation of IUPM^38^.

For P1, blood monocytes were isolated before the initiation of ART (viral load of 31,800 copies/mL) and four months after initiation of ART (when circulating virus was undetectable) and QVOAs were performed (**Table 2**). For P1, viral RNA was detected in the MC QVOAs for both the pre- and post-treatment samples, and the IUPM fell from 1.1 to 0.24 over the treatment period. For P2, QVOAs were performed on AM, MC and CD4^+^ T cells, but only the AM and T cells were positive for viral RNA (IUPM in the range 0.5 to 0.8). For P3, viral RNA was detected in MC (3.7 IUPM) and CD4^+^ T cells (0.66 IUPM), but not in AM. For P4 and P5, QVOAs were performed on MC and CD4^+^ T cells and viral RNA was detected in both (**Table 2**) (**Fig 4b**). As expected, these numbers of productively infected peripheral blood MC and T cells are much less than the mean levels of viral DNA previously detected using ddPCR in these cell types isolated from 6 ART-suppressed donors (∼700 copies/10^6^ T cells and ∼60 copies/10^6^ MC)^20^. It is commonly accepted that most viral DNA copies are not replication competent, and further, the QVOA assay is not 100% efficient. Thus, these results provide a minimum estimate of the level of productively infected HIV target cells^39^.

### Infectivity of viruses obtained from QVOA supernatants

To further establish that the viral particles present in the AM-QVOA, MC-QVOA and T cell QVOA supernatants contained replication competent virus, we spin-inoculated MOLT-4/CCR5 target cells and followed the time course for appearance of nascent viral RNA in the culture supernatants using RT-qPCR **(Fig. 5a).** Supernatants were collected at days 0, 4, 10, and 15 after spin-inoculation and viral spread was observed in all MC and T cell QVOAs for P2, P4 and P5 (**Fig. 5b, 5d, 5e**). However, for P3 viral spread was only observed in MC, not T cells (**Fig. 5c)**. We estimated that 0.7 % to 2.2 % of the viral particles present in the MC and T cell QVOAs were replication competent. We also calculated an upper limit of 0.2 % for replication competent virus present in the P3 T cell QVOA supernatants based on the absence of outgrowth (**Fig. 5f**).

**Figure 5.**
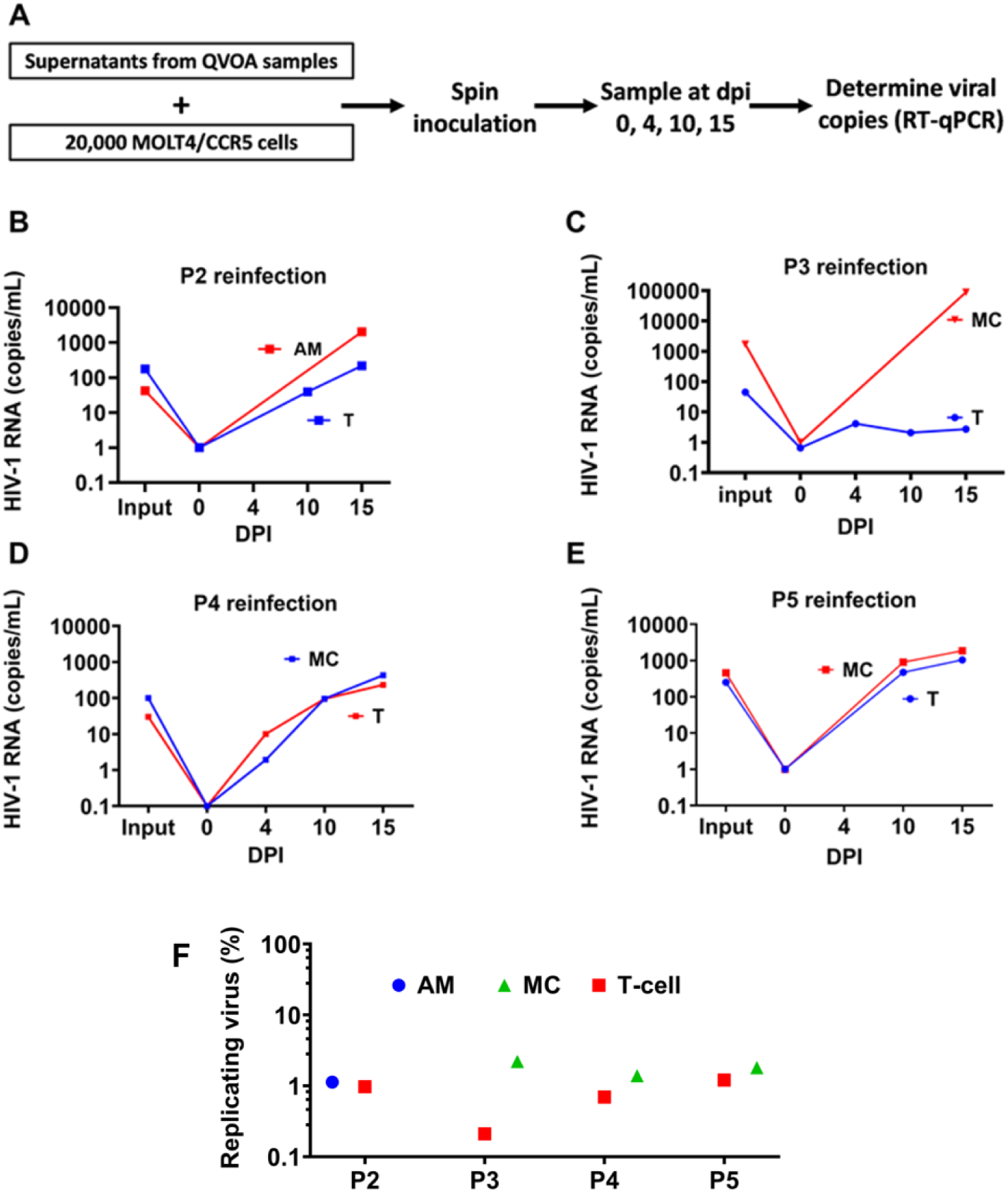
Viruses collected from QVOA supernatants of AM, MC and T cells can establish *de novo* infection. (**A**) Supernatants from QVOA positive wells were used to separately spin infect 200,000 MOLT4/CCR5 cells using serial dilutions. Supernatants were collected at days 0, 4, 10, and 15 after spin infection and HIV RNA copies were determined using RT-qPCR. (**B**) Reinfection kinetics of AM and T cells obtained from P2 QVOA supernatant. (**C**) Reinfection kinetics of MC and T cells obtained from P3 QVOA supernatant. (**D**) Reinfection kinetics of MC and T cells obtained from P4 QVOA supernatant. (**E**) Reinfection kinetics of MC and T cells obtained from P5 QVOA supernatant. (**F**) Percentage of replicating viral particles in HIV RNA copies detected in QVOA. HIV RNA copies in the culture supernatants were measured by RT-qPCR and the frequency of replicating viral particles was calculated using an Infection Frequency Calculator developed using limiting dilution statistics based on the number of positive wells and the input number of HIV copies (https://silicianolab.johnshopkins.edu). The replicating virus percentage of AM, MC, and T cells for each patient assayed is shown. The data in B, C, D, and E show the mean value of three technical replicates.

### Major viral variants isolated from AM, MC and T cell QVOAs

We used targeted amplicon next generation sequencing to perform a sequence analysis of the V3 regions of HIV env using virus isolated from the QVOA wells from CD4^+^ T cells, AM and MC from each patient (**Fig. 6a, 6b**). Comparisons of the predominant viral sequences isolated from myeloid cells (AM or MC) and T cell QVOAs from the same donor showed 100 % (P2), 100 % (P3), 91 % (P4), and 97 % (P5) pairwise identity. Further, all major variants analyzed were predicated to use the R5 co-receptor (**Supplemental Table S1**). These data, which only reliably report on the major variants present, do not address the question of whether common or distinct myeloid and T cell viral pools exist. The predicted amino acid sequences of each major variant were matched to a single HIV consensus sequence calculated from 2635 HIV clinical sequences archived in the Los Alamos HIV sequence database^40^. Amino acid substitutions were clustered in various regions with the same pattern for P2 and P3 and more variabilities for P4 and P5 (**Fig. 6c**). An APOBEC-induced hypermutation analysis using Hypermut 2.0 (https://www.hiv.lanl.gov/content/sequence/HYPERMUT/hypermut.html) did not reveal any definitive A3G or A3F deamination sites (**Supplemental Table S2**).

**Figure 6.**
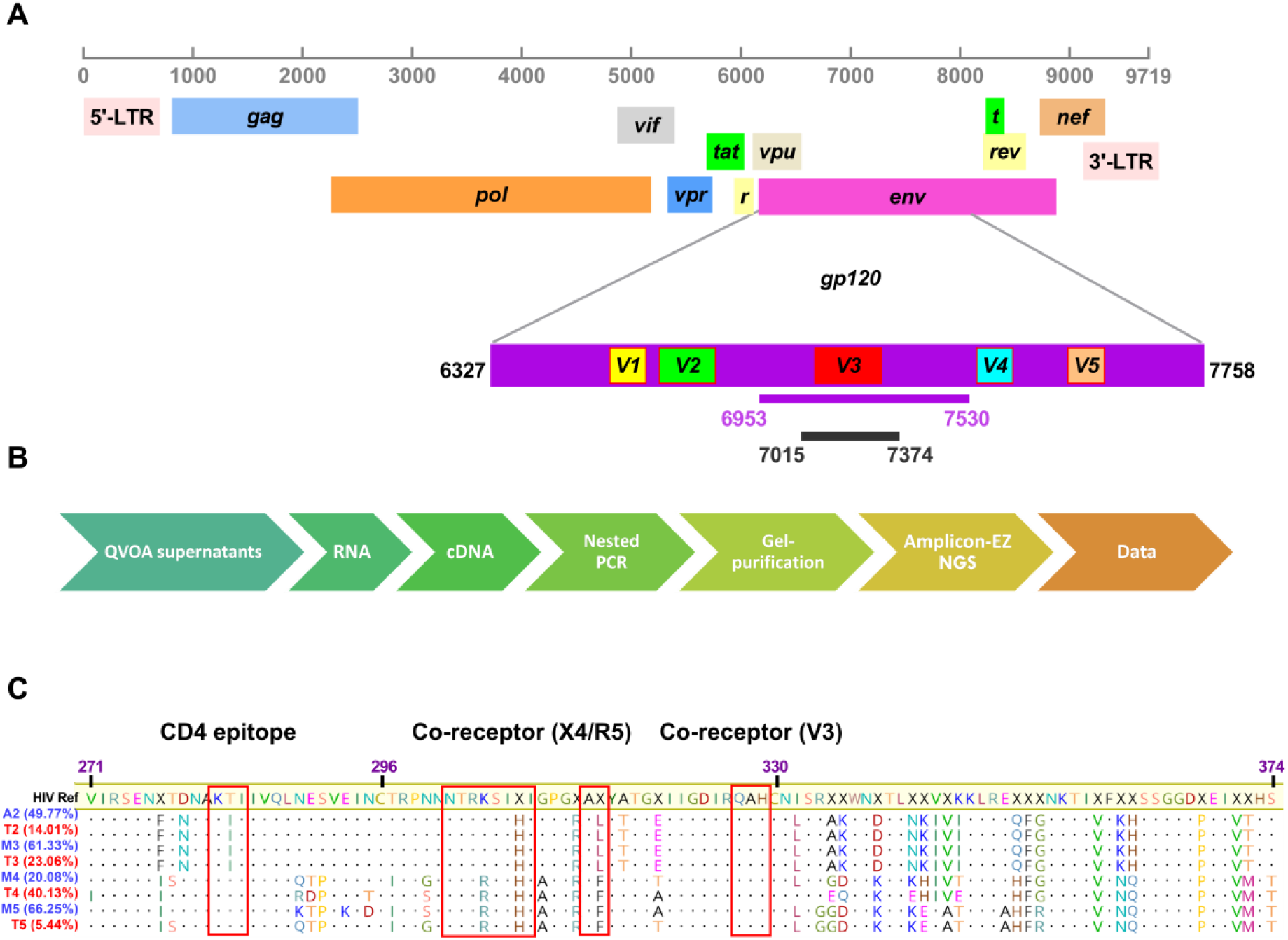
Sequence analyses of virus produced in MC, AM, and CD4^+^ T cell QVOA. (**A**) The HIV-1 genome structure and the region targeted for sequencing is shown. The first PCR (6953-7530) and second PCR (7015-7374) regions were targeted by nested PCR primers using cDNA obtained from AM, MC, and T cell QVOA’s from P2 to P5. The purified amplicons (359 bp) were then sequenced using Illumina MiSeq. (**B**) Outline of the stepwise procedure used in targeted amplicon sequencing. (**C**) Major env amino acid sequences derived from sequencing extracellular viral RNAs produced in QVOAs. The consensus reference sequence is derived from 2635 HIV-1 clinical isolates. Boxed regions at the protein sequence level show the mutation spectrum within the CD4-associated and co-receptor binding sites. The positions are relative to the HXB3 strain. Sequences derived from myeloid cell infections are indicated in blue [either monocyte (M) or alveolar macrophage (AM)] and sequences obtained from T cell infections are in red with the numerical patient identification shown in Tables 1 and 2. The major variant percentages are indicated.

## DISCUSSION

Macrophages exhibit malleable and temporal phenotypes *in vivo* determined by their microenvironment and cytokine stimuli. Given this property, it is not possible to evaluate the “average” behavior of a macrophage with respect to an experimental variable such as HIV infection despite the isolation of cells from a defined immune compartment. Even *in vitro* experiments using highly purified monocyte derived macrophages (MDM) and defined strains of virus are complicated by the presence of a mixed population of cells with respect to their inflammatory phenotype, cell cycle markers (i.e. G0 or G1) and other variables. No single study using primary cells can evaluate all the variables that may impact the macrophage phenotype and susceptibility to HIV infection. In this work, *in vivo* HIV infection of monocytes (MC) and alveolar macrophages (AM) is explored with a focus on whether previous *in vitro* findings concerning the nucleotide pool composition of MDM and the involvement of uracil base excision repair (UBER) extend to *in vivo* infections. These studies also provide rare (but limited) measurements of the replication capacity of proviruses present in circulating MC and resident alveolar macrophages of the lung.

### Implications of AM and MC dNTP pool composition and UBER status on *in vivo* infection

As noted earlier, three well-documented hallmarks of *in vitro* infection of MDM and MC are low dNTP pools, the presence of elevated dUTP, and the incorporation of dUMP into viral reverse transcripts. We have previously established using *in vitro* systems that the outcome of HIV infection in the presence of high dUTP pools will critically depend on the expression level of the UNG enzyme^20, 30, 31^. If its expression is relatively high, uracilated viral DNA products are degraded before integration and only non-uracilated viral DNA is integrated^31^. If its expression is low or absent, uracilated viral DNA can integrate but the expression of viral RNA transcripts and protein is decreased^20, 30, 31^. Our previous uracil excision ddPCR measurements detected high levels of viral uracilation in MC samples from six HIV-infected donors and AM samples from a single donor, strongly suggesting that the features of elevated dUTP/dTTP levels and viral incorporation of dUMP that have been documented for *in vitro* infections of MDM and MC extend to *in vivo* infections of AM and MC. This conclusion is directly validated in this work where we have directly determined both low overall dNTP pool levels in AM and elevated dUTP. Further, our findings that UBER enzyme expression is severely compromised in AM, and to a lesser extent MC, provides an explanation for both the detection and persistence of viral uracils in the previous uracil excision ddPCR study. Ironically, the potential for a strongly restrictive effect of high dUTP is blunted by the absence of UNG activity in AM. Accordingly, the major outcome of infection under these conditions is persistence of uracilated proviruses (ie. for the lifetime of the AM, which is estimated at several months)^41^. Given our previous finding that RNA pol II transcription from uracilated DNA templates is downregulated compared to templates containing only T/A^42^, the uracilated proviruses are expected to be at least partially repressed in terms of transcription. Since U/A pairs have been reported to have significant effects on protein-DNA interactions (usually disruptive)^35, 41–44^, it is possible that alterations in the chromatin structure may be unfavorable for transcription or that binding of certain required HIV transcription factors may be impaired. Elaboration of these effects is the subject of future investigations, but the currently available data suggests that abundant U/A pairs in proviruses would tend to be transcriptionally repressive and may even contribute to a unique mechanism of latency in long-lived macrophages such as microglial cells of the brain.

### Implications of replication competent viral sequences in AM and MC

An important finding in this study is that rare replication competent viruses can be detected in both AM and peripheral MC from HIV-infected individuals that were virally suppressed using ART. Beyond the obvious need to control for contaminating T cells in AM and MC QVOAs, a frequent critique is that phagocytized infected T cells containing integrated virus could give rise to the observed signals. A phagocytic mechanism is difficult to discount with one hundred percent certainty, but is made unlikely given that on average ∼80% of infected MC and AM cells contain uracilated viral DNA and we previously established that viral uracilation is a unique phenotype of myeloid cell infection (i.e. integrated virus from a phagocytized T cell would not contain dUMP)^20, 31^. The frequencies of replication competent viruses that we observed with peripheral MC overlap with findings from previous studies using human and macaque samples, and detection of rare replication competent virus in human AM samples has also been reported (comprehensively reviewed in Wong *et al*)^2^. Although our sample number is low for AM due to halting of bronchiole alveolar lavage procedures during the Covid-19 pandemic, the limited data suggest that the frequency of replication competent virus may be smaller in this compartment compared to peripheral MC.

The presence of replication competent virus in both MC and AM when the lifetimes of these cells *in vivo* is in the time frame of several days and months, respectively, strongly suggests that these cells are encountering active virus during their lifetimes. For the MC population, new findings on the biology of these cells indicates that they can frequently migrate through immune cell compartments without differentiation into tissue macrophages^45^. This migration pattern increases the chances that these cells will enter a sanctuary compartment containing infected CD4^+^ T cells or macrophages, become infected, and then reenter circulation. In contrast, AM are stationary within the lung tissue and must encounter virus through infected neighbor cells or through transient encounter with rare infected CD4^+^ T cells. For these active replication mechanisms to be viable, sanctuary compartments where antiretroviral drugs are poorly penetrant are required. In addition, macrophages are more resistant to many antiretroviral drugs and are less prone to cell death after HIV infection compared to T cells, which increases the time frame for release of virus particles from these cells^46, 47^.

## CONCLUSIONS

This study adds to the growing body of evidence that myeloid lineage cells can harbor rare replication competent HIV after infected patients have undergone ART and achieved viral suppression. A biomarker for infection of these cells is viral DNA that contains dUMP.

Since the predominant G0 population of macrophages has vastly diminished UBER activity, the fate of viral dUMP is limited to persistence and potential inhibitory effects on transcription based on *in vitro* studies. For short lived macrophage and monocyte cells, the persistence may last for the lifetime of the cells. For longer lived brain microglial cells, the opportunity increases for cells to enter a transient G1 state which would allow the excision repair process to proceed and potentially repair the U/A pairs, decrease transcriptional repression, and potentially lead to viral-induced inflammatory conditions often observed in the brains of HIV infected patients.

## METHODS

### Study subjects

This study was approved by the Johns Hopkins Institutional Review Board and informed consent was obtained from all participants before study enrollment. Peripheral blood was obtained from healthy volunteers and ART suppressed patients with viral loads <20 copies HIV-1 RNA/mL. Monocytes and resting CD4^+^ T cells were obtained from donor PBMCs as described below. Characteristics of study participants are given in **Table 1**. Collection of BAL samples were discontinued in March 2020 due to the extreme risks pertaining to the Covid-19 pandemic. Therefore, sample numbers are unavoidably reduced from the original planned enrollment for the study.

### Isolation of primary cells from donor PBMCs

Monocytes (MC) were purified from peripheral blood mononuclear cells (PBMC) using a Ficoll-Hypaque density gradient followed by negative selection using a Pan monocyte isolation kit (Miltenyi Biotech). Then MCs were differentiated into monocyte-derived macrophages (MDM) by culturing MC under adherent conditions for seven days using MDM-AS medium containing RPMI 1640 (Gibco) supplemented with 10% donor autologous serum, 1% Pen/Strep (HyClone), 0.3 mg/mL glutamine and 10 mM HEPES, plus 10 ng/mL M-CSF (R&D Systems). Fully differentiated MDMs were maintained in media containing RPMI 1640 plus 10% dialyzed FBS, 1% Pen/Strep.

CD4^+^ T cells were isolated from PBMC population using the CD4^+^ isolation kit (CD4^+^ T cell Isolation kit II, Miltenyi Biotec, San Diego, CA). CD4^+^ T cells were first cultured for three days using stimulating conditions (RPMI supplemented with 10% donor autologous serum, 1% Pen/Strep (HyClone), 0.3 mg/mL glutamine, 1 mg/mL of Phytohemagglutinin (Gibco) and, ciprofloxacin (5 mg/mL)).

The dividing cell line HAP1 (Horizon Discovery) was seeded at 2 million cells per T-75 flask. The culture medium containing RPMI-1640 medium plus 10% heat inactivated FBS, 1% Pen/Strep. Cells were grown until 70% confluence with medium changes every 48 h. The cells were released from the flask using 5 mL of 0.25% trypsin-EDTA (Gibco), followed by washing three times with PBS (Gibco).

### Processing of alveolar macrophages (AM) from bronchoalveolar lavage procedure (BAL)

Bronchoscopy and broncho-alveolar lavage (BAL) were performed on six participants including three healthy donors, a single HIV-1 infected patient pre- and post-ART treatment and two HIV-1 patients on ART (**Table 1**). To obtain alveolar macrophages (AM), bronchoscopy and lavage were done, as described elsewhere^48^. Briefly, BAL samples were filtered using a sterilized gauze pad and transferred into 50-mL centrifuge tubes. A pellet was obtained using a short spin (250 × *g* for 5 min) and was resuspended in serum-free RPMI 1640 medium (Gibco) containing antibiotics (Pen/Strep, gentamicin, and amphotericin B) and incubated at 37°C for 24 h without shaking. Following incubation, the cells were washed and resuspended in culture medium containing RPMI-1640 medium plus 10% heat inactivated FBS, 1% Pen/Strep. Cells were then counted and diluted to 10^6^ cells/mL for further use. Cells were then pelleted and used for dNTP extraction, or *in vitro* infection, or quantitative viral outgrowth assay (QVOA) as described below.

### Generation of RNA standards for TCR β RNA assay

A fragment of TCR β containing both V-region and C-region was subcloned into a mammalian expression vector. The plasmid was transformed to DH5α competent cells then spread onto an agar plate containing corresponding antibiotics. A mini prep was performed followed by plasmid purification and quantification. Two sets of real time PCR primers targeting both V-region (68bp) (rt-FWD 5’-ACACGTGAAATGCTCTTTGCG-3’, rt-REV 5’-TTACTCCTGCGCCTCTGTGTC-3’) and C-region (65bp) (rt-FWD 5’-TGGCTTCTGGCACTCCTTG-3’, rt-REV 5’-GCCATGTGAAGACAGAGGCA -3’) were used to generate standard curve.

### Quantitation of TCR β RNA

AM and MDM purity was checked for T-cell contamination by RT-qPCR using TCR-β primers. TCR β RNA was quantitated by RT-qPCR using the above-described primers. Cycling conditions were as follows: 95 °C for 5 min, and 40 cycles of 94 °C for 10 s, 55 °C for 15 s, and 60 °C for 30 s.18S rRNA was simultaneously detected with the TCR β RNA to control for cell counts. To determine the average number of TCR β copies, AMs and MDMs from three donors, were isolated and purified using the above-described methods. The purity of the cells was confirmed by marker staining, and the relative number of copies of TCR β RNA was compared to CD4^+^ T cells.

### Characterization of isolated cell populations

The function and cell identity of AM, MDM, and CD4^+^ T cells was determined by morphology, marker staining, functional assays, and RT-qPCR (see above). The purity of AM cells was analyzed by marker staining using anti-CD68 antibody ([EPR20545] (ab213363), abcam) and pHrodo™ Green E. coli BioParticles™ Conjugate for Phagocytosis (P35366, Thermofisher). Using this method, we obtained AMs with a purity of >95.8% and are functional of >91.6% with neglectable T cell contamination (0.9% for AMs and 1.6% for MDMs).

### Single nucleotide polymerase extension assay

The extraction of total dNTPs from ∼ 1-2 million cells and quantification of dUTP and dTTP were performed as previously described^30, 31, 49^, except that HIV-1 reverse transcriptase (RTase) was used in the extension assay (Millipore-Sigma). The extension reactions were performed at 37 °C for 40 min and included ∼6 to 1200 fmol of total dNTP, 120 or 1200 fmol of DNA template-primer for extracts obtained from non-dividing or dividing cells, respectively, and 2 units of RTase (one unit converts 1 nmol of labeled dTTP into acid-insoluble material in 20 min at 37°C, pH 8.3). The substrate (22 mer) and the product (23mer) of the extension reaction were separated by electrophoresis on a denaturing PAGE (8 M urea, 1× TBE, 15 % acrylamide) and then imaged (Typhoon 9500 imager; GE Healthcare) and the images were quantified using Image Lab software (Bio-Rad). The extension buffer consisted of 20 mM Tris-HCl, pH 8.0, 100 mM KCl, 5 mM MgCl_2_, 2 mM DTT, and 0.1% BSA. Extensive validation experiments were performed using [dUTP + dTTP] mixtures with known dUTP/dTTP 1:1 ratio to establish a detection limit and a linear correlation between the known dNTP concentrations and the ratio determined from the SNE assay. The amount of dUTP per million cells was determined by first calculating the pmol dUTP in the SNE reaction = [pmol DNA template-primer] [fractional extension (no dUTPase) – fractional extension (add dUTPase)]. The total pmol dUTP in the methanol extract was calculated after correction for dilutions and the fraction of the total extract analyzed in the SNE reaction. The total pmol of dUTP was then divided by the cell number used to prepare the extract. The cell number was normalized to the genomic RPP30 copies as the standard (diploid for AM and MDM, and haploid for HAP1).

### Expression levels of UBER and dNTP metabolic enzymes using real-time qPCR

RNA was extracted from AM, MDM or HAP1 cells using the RNeasy Mini Kit (Qiagen) following the manufacturer’s protocol. About 500 ng of RNA was used to synthesize cDNA with High-Capacity cDNA Reverse Transcription Kit (Applied Biosystems, #4368813) following the manufacturers protocol. The cDNA was used as input material and the Rotor Gene qPCR probe kit (Qiagen, #204374), mRNA copies of infected AM and MDM or uninfected controls were measured and normalized to an internal control gene (18S rRNA). Thermal cycling conditions for qPCR consisted of 95 °C for 5 min, and 40 cycles of denaturation at 95 °C for 10 sec and annealing and extension at 60 °C for 30 sec. The relative expression levels were calculated using the 2^-ΔΔCT^ method, where the ΔΔC_t_ is the difference in ΔC_t_ between the target and reference samples:

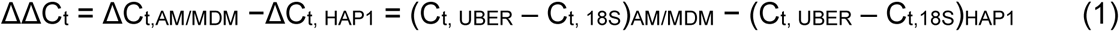

### Viruses and infections

The macrophage tropic replicative HIV^Bal^ was obtained from the NIH AIDS Reagent Program (NIH-ARP #11414) and propagated in MOLT-4/CCR5 cells. Culture supernatants of infected MOLT-4/CCR5 cells were collected at 14 dpi, centrifuged to clear cellular debris, passed through 0.45 μm filters, aliquoted, and stored at -80 °C. Viral titer was determined by the ELISA p24 antigen assay (Lenti-X p24 Rapid Titer Kit, TaKaRa). Infections of fully differentiated MDM were performed after adherent growth in the presence of M-CSF for seven days. Just prior to addition of virus to MDM, the media was replaced with RPMI + 10% dFCS and omitting M-CSF. Infections of purified AM were performed after adherent purification. AMs were seeded at 1 x 10^5^ cells per well in a 24-well plate and kept at 37 °C, 5% CO2. The culturing medium was removed, and cells were washed twice with PBS, then infected with HIV^Bal^ strain at the multiplicity of infection (MOI) of 1. Then the cells were washed with PBS and fresh medium was replaced.

### Determination of viral copy number using real-time qPCR

Viral copies were determined using qPCR. A standard curve was generated using 10-fold serial dilutions of DNA extracted from J-lat cells (NIH-ARP). Real-time qPCR was performed using Rotor-Gene Probe PCR kit (Qiagen, #204374) on Rotor-Gene qPCR instrument. PCR targeted the early or late HIV reverse transcripts (ERT or LRT). Amplification was performed using a two-step program: initial heating at 95 °C for 5 min, followed by 40 cycles of denaturation at 95 °C for 10 sec and annealing and extension at 60 °C for 30 sec. For determination of viral copies per cell, simultaneous quantification of RPP30 gene was performed using published primers and a specific probe as previously described. All primers and probes used in this study are listed in **Supplemental Table S3**.

### Determination of HIV proviral DNA using *Alu-gag* nested PCR

To determine copy numbers of HIV proviral DNA, a nested PCR method was used. The first PCR was performed using a forward primer (Alu-F, 0.2 μM) that targeted genomic *alu* sequences randomly located near integrated proviruses and an HIV-specific *gag* reverse primer (Gag-R, 1.2 μM). Other PCR conditions were 200 μM of each dNTP, 1X LongAmp taq buffer (NEB), 5 units of LongAmp taq DNA polymerase (NEB) and 50 ng of DNA sample extracted from infected cells in a 50 μL final reaction volume. Amplification was performed using the following thermocycler program: initial activation heating 94 °C for 2 min, followed by 20 cycles of denaturation at 94 °C for 30 sec, annealing at 50 °C for 30 sec and extension at 65 °C for 3 min and a final extension reaction at 65 °C for 10 minutes. The PCR product is diluted 20-fold and five μL of the diluted PCR product is used as an input material for the second PCR reaction, which is performed using LRT forward and reverse primers and probe using the Rotor-Gene Probe PCR kit (Qiagen) as described above. Proviral copy numbers were determined using the J-lat cell integration standard as previously described^20, 30, 31^. Genomic DNA copy numbers were determined using the RPP30 reference gene using the same amount of input DNA used to measure proviral copy numbers.

### Uracil content of viral DNA

Uracil content of viral DNA was determined using uracil excision qPCR method (Ex-qPCR) as previously described with some modifications as described below^20, 30, 31^. Ex-qPCR was used to determine uracil-containing fraction of viral DNA. The sample was split into two equal portions and one portion was treated with UDG. Briefly, 0.2 units of UDG (NEB) was added into the qPCR master mix to excise uracils from viral DNA. The qPCR reaction was modified to include the UDG reaction time and heat-cleavage of the resulting abasic sites. Thermocycler program for this reaction was: 37 °C for 30 min (UDG reaction), 95 °C for 5 min (abasic site cleavage) and 40 cycles of denaturation at 95 °C for 10 sec and annealing and extension at 60 °C for 30 sec. Primers and probe sets used to amplify viral DNA are indicated under the specific experiments listed in **Supplemental Table S3**. Primers and probe sets targeting RPP30 were used to calculate Frac U^DNA^ using the ΔΔC_t_ method.

### CD4^+^ T cells viral outgrowth assays (CD4-QVOA)

Peripheral blood mononuclear cells (PBMCs) from healthy HIV-1-infected patients (under suppressive cART with undetectable plasma viral loads) were separated from whole blood by a standard Ficoll gradient. CD4^+^ T cells were obtained using a negative CD4 magnetic selection kit (Stemcell Technologies), according to the manufacturer’s instructions. To obtain activated T-cell blasts, the cells were resuspended in Roswell Park Memorial Institute (RPMI) medium (RPMI 1640 medium, 10% fetal bovine serum, 2 mM L-glutamine, 100 U mL^−1^ penicillin and 100 μg mL^−1^ streptomycin (Invitrogen)) and treated with 5 μg mL^−1^ PHA (Sigma–Aldrich) and 10 U mL^−1^ IL-2 (Roche) for 48 h at 37 °C. The cells were then re-suspended in RPMI medium, seeded in successive 1:5 or 1:10 serial dilutions in triplicate wells of a 6-well plate or 24-well plate (starting from 1 × 10^6^ cells), and reactivated for 48 h at 37 °C with 5 μg mL^−1^ PHA. Next, the cells were washed, resuspended in fresh medium containing 10 U mL^−1^ IL-2, and co-cultured with MOLT-4/CCR5 cells (0.5 × 10^6^ cells) for 2 weeks at 37 °C, with fresh medium containing IL-2 replenished after 1 week. HIV-1 in the co-culture supernatants was measured using RT-qPCR. Infection frequencies were determined by maximum likelihood statistics as described previously, using the application at http://silicianolab.johnshopkins.edu, and expressed as infectious units per million (IUPM).

### AM viral outgrowth assays (AM-QVOA)

BAL samples were collected from HIV-1 infected patients and AMs were isolated and purified as described above. The cells were then re-suspended in RPMI medium, seeded in successive 1:5 or 1:10 serial dilutions in triplicate wells of a 6-well plate or 24-well plate (starting from 1 × 10^6^ cells), and kept for 72 h at 37 °C with T20 entry inhibitor (10 μM) to prevent reinfection. Next, the cells were washed and co-cultured with MOLT-4/CCR5 cells (0.5 × 10^6^ cells) for 2 weeks at 37 °C, with fresh medium containing lipopolysaccharide (Sigma-Aldrich, L5293) replenished after 1 week. HIV-1 in the co-culture supernatants was measured using RT-qPCR. Infection frequencies were determined by maximum likelihood statistics as described previously, using the application available online at http://silicianolab.johnshopkins.edu, and expressed as infectious units per million (IUPM).

### MC viral outgrowth assays (MC-QVOA)

Blood samples were collected from HIV-1 infected patients and MCs were purified and differentiated with T20 entry inhibitor (10 μM) to prevent reinfection as described above. The cells were then re-suspended in RPMI medium, seeded in successive 1:5 or 1:10 serial dilutions in triplicate wells of a 6-well plate or 24-well plate (starting from 1 × 10^6^ cells) and co-cultured with MOLT-4/CCR5 cells (0.5 × 106 cells) for two weeks at 37 °C, with fresh medium containing lipopolysaccharide (Sigma-Aldrich, L5293) replenished after 1 week. HIV-1 in the co-culture supernatants was measured using RT-qPCR and IUPM was calculated as above.

### HIV-1 reinfection assay

MOLT-4/CCR5 cells (20,000/well) were subjected to spin-inoculation for 2 h with diluted supernatant containing varying copies of HIV-1 RNA from positive QVOA wells. Supernatants were collected at days 0, 4, 10, and 15 after spin-inoculation. RNA was isolated from supernatant, and HIV-1 RNA was quantitated by RT-qPCR and number of replicating HIV-1 particles was calculated by maximum likelihood statistics as described above using the application available online at http://silicianolab.johnshopkins.edu.

### Next-generation Amplicon-EZ sequencing of HIV RNA

HIV-1 RNA from the QVOA supernatants was isolated using PureLink™ Viral RNA/DNA Mini Kit (Invitrogen, #12280050) and RT-PCR was performed using the High-Capacity cDNA Reverse Transcription Kit (Applied Biosystems, #4368813) following the manufacturers protocol. After cDNA synthesis, a 359 bp segment of the env region was amplified using nested PCR (outer primers E90 and Nesty8 and inner primers DLoop and E115, as described previously)^50^. Thermocycler settings were the same for both outer and inner env PCR reactions: 94 °C for 3 min, followed by 35 cycles and 40 cycles of: 94 °C for 30 s, 55 °C for 30 s and then 68 °C for 1 min, followed by a final extension at 68 °C for 5 min for the first and second PCR, respectively. The PCR products were gel extracted using the QIAquick Gel Extraction Kit (Qiagen, # 28706) and quantified using a Nanodrop 2000. The amplification product (20 ng/uL, 50 uL total) were sent to GENEWIZ for targeted amplicon sequencing. Sequencing adapters are listed in **Supplemental Table S4.**

DNA library preparations were performed using NEBNext Ultra DNA Library Prep kit following the manufacturer’s recommendations (Illumina, San Diego, CA, USA). Briefly, end repaired adapters were ligated after adenylation of the 3’ends followed by enrichment by limited cycle PCR. DNA libraries were validated and quantified before loading. The pooled DNA libraries were loaded on the Illumina instrument according to manufacturer’s instructions. The samples were sequenced using a 2 x 250 paired-end (PE) configuration. Image analysis and base calling were conducted by the Illumina Control Software on the Illumina instrument.

Unique sequence and abundance raw sequence data were demultiplexed using bcl2fastq version 2.17.1.14. Read pairs were trimmed for adapter sequences and low quality basecalls using Trimmomatic version 0.36. Reads were discarded if they were less than 30 bases long. Each read pair was then merged using the bbmerge tool from the bbtools software toolkit. Reads that could not be merged were discarded from further analysis. The target sequence between conserved flanking primers was extracted from each merged pair. The forward and reverse reads were joined and assigned to samples based on barcode and truncated by cutting off the barcode and primer sequence. Quality filtering on joined sequences was performed (**Supplemental Table S4**). The sequences are grouped according to their abundance and identities (single nucleotide differences were considered different).

### Determination of viral tropism

Viral tropism was predicted according to the V3 region of the most abundant sequence of each QVOA by applying the 11/25 rule with a false-positive rate of 10%^51^, and the Position-Specific Scoring Matrix (PSSM) approach (https://indra.mullins.microbiol.washington.edu/webpssm) **(Supplemental Table S1)**^52^**. Fluorescence microscopy.** AMs or MDMs were treated at 37°C for 4 h with 10 μM pHrodo Green *E. coli* BioParticles (Invitrogen, P35366), which can be phagocytosed only by functional macrophages and are nonfluorescent at neutral pH, but which are fluorescent in the acidic pH of phagosomes. The cells were fixed using cold methanol for 15 min then stained at room temperature for 60 min with anti-CD68 antibodies and DAPI for 5 min. Images were taken on an Olympus IX81 fluorescence microscope and analyzed using Fiji ImageJ.

### Statistics

The data uncertainties are expressed as the mean ± 1 SD. Unless otherwise indicated in the figure legends, data are derived from three independent biological replicates using cells isolated from three donors. Experiments using cells from individual donors were performed in technical triplicates. Data were analyzed using the Prism 8.0 statistical program from GraphPad Software. Pairwise statistical analysis of test groups was performed by a two-tailed unpaired Student’s *t*-test, assuming equal variances between groups. Due to the Covid pandemic, all BAL clinical procedures were prohibited in March 2020 which prevented collection of the entire planned set of samples. Due to this unanticipated event, the uncertainty in some of the measurements with patient samples is unavoidably greater.

## ABREVIATIONS

AM: alveolar macrophage
APE1: Apurinic/apyrimidinic endonuclease 1
A3A: Apolipoprotein B mRNA editing enzyme, catalytic polypeptide-like 3A
A3G: Apolipoprotein B mRNA editing enzyme, catalytic polypeptide-like 3G
ART: antiretroviral therapy
BAL: Bronchoalveolar lavage
BLT: bone marrow-liver-thymus
cDNA: Complementary DNA
CNS: central nervous system
DAPI: 4’,6-diamidino-2-phenylindole
ddPCR: Droplet Digital PCR
dNTP: Deoxynucleoside triphosphate
dsDNA: double stranded DNA
dU: deoxyuridine
dUMP: deoxyuridine monophosphate
DUT: DUTP pyrophosphohydrolase
dUTP: deoxyuridine triphosphate
dT: Thymidine
dTTP: Thymidine triphosphate
Env: viral envelope protein
ERT: early reverse transcripts
HAP1: a near-haploid human cell line
HIV: human immunodeficiency virus
hUNG: human uracil DNA glycosylase
IDT: Integrated DNA Technologies
IUPM: infectious units per million
Lig III: DNA Ligase 3
LRT: late reverse transcripts
LTR: long terminal repeat
MC: monocyte
M-CSF: Macrophage colony-stimulating factor
MDM: Monocyte-derived macrophage
NEB: New England Biolabs
P24: a component of the HIV particle capsid
PBMC: peripheral blood mononuclear cell
PHA: Phytohemagglutinin
Pol β: DNA polymerase beta
QVOA: Quantitative Viral Outgrowth Assay
RPP30: Ribonuclease P/MRP Subunit P30
RT-PCR: Reverse transcription polymerase chain reaction
SAMHD1: SAM domain and HD domain-containing protein 1
SIV: simian immunodeficiency virus
TCR β: T-cell receptor chain beta
TMP: Thymidine monophosphate
TTP: Thymidine triphosphate
UBER: uracil base excision repair
UNG: uracil DNA glycosylase

## DECLARATIONS

### Ethics approval and consent to participate

The protocols described in this work have been approved by the Johns Hopkins University Internal Review Board (IRB 00038590) and follow established ethical practices with informed donor consent.

### Consent for publication

Not applicable

### Availability of data and materials

The sequence datasets generated and/or analyzed during the current study are available in the GenBank repository: Sequence Read Archive (SRA); BioProject ID PRJNA832816. Supporting data generated or analyzed during this study are included in this published article and its supplementary information files. The complete data for this study is accessible for download without any restrictions at: [5-03-2022_CUI_AM_Ms_DATA](https://livejohnshopkins-my.sharepoint.com/:f:/g/personal/jstiver1_jh_edu/Ev081-qXPgFKufidlaQvw0wB0LRcBGlu3kr5kcRQcx5Vxg?e=NrK2tP)

### Competing interests

The authors declare that they have no competing interests.

### Funding

This work was supported by NIH grants R01 AI124777 and RO1 GM056834.

### Authors’ contributions

JC: Conceptualization, Data curation, Formal analysis, Investigation, Methodology, Validation, Writing – original draft, Writing – review & editing

MM: Formal analysis, Investigation, Methodology, Validation NC: Formal analysis, Methodology

CM: Performed bronchial alveolar lavage procedures

EF: Prepared IRB protocols, coordinated all clinical procedures

JB: Prepared IRB protocols, coordinated all clinical procedures, assisted in clinical procedures

JJ: Approved donor consents, assisted in recruitment of HIV patients

JTS: Conceptualization, Formal analysis, Funding acquisition, Methodology, Project administration, Resources, Supervision, Validation, Visualization, Writing – original draft Writing – review & editing

## Acknowledgements

The authors acknowledge the support of the Johns Hopkins Institute for Clinical and Translational Research team and especially Ms. Christina Bunch for recruiting, consenting, and performing blood draws from healthy subjects. The authors thank Dr. Rebecca Veenhuis (Johns Hopkins School of Medicine) for QVOA protocol optimization.

## Notes

https://livejohnshopkins-my.sharepoint.com/:f:/g/personal/jstiver1_jh_edu/Ev081-qXPgFKufidlaQvw0wB0LRcBGlu3kr5kcRQcx5Vxg?e=NrK2tP

### Competing Interest Statement

The authors have declared no competing interest.

