## Supplemental Information for "Replication-competent HIV-1 in human alveolar macrophages and monocytes despite nucleotide pools with elevated dUTP"

**for**

<u>TABLE OF CONTENTS</u>

FIGURE S1

FIGURE S2

SUPPLEMENTAL TABLES S1-S4

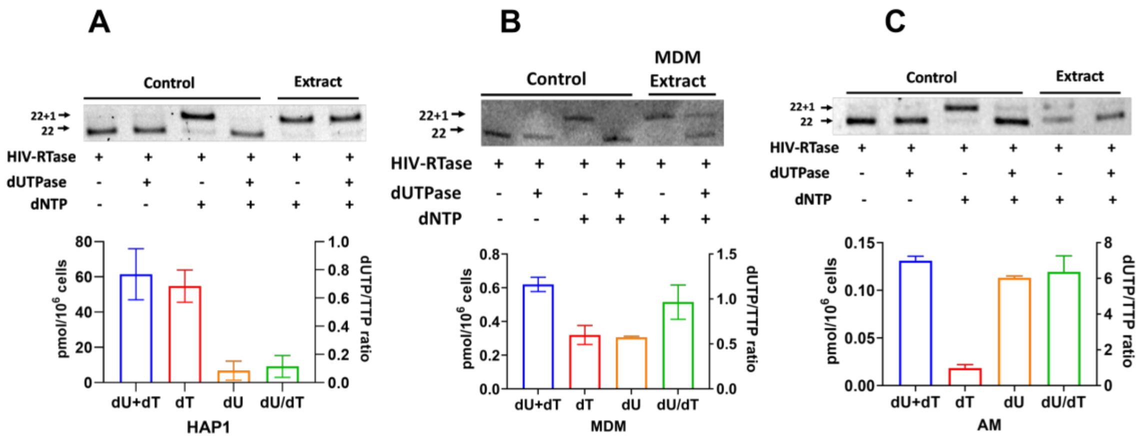


**Figure S1. Characterization of dUTP and dTTP levels in HAP1, AM, and MDM cells.** The single nucleotide extension assay was used to establish the differences in dUTP/dTTP between MDM, AM and HAP1 dividing cells. The procedure is described in Methods. **(A)** (top) Denaturing urea-PAGE of extension reactions in the presence and absence of dUTPase with cell extract from HAP1 cells. The image is from one of three biological replicate measurements. (bottom) Total levels of [dTTP + dUTP], dTTP alone, dUTP alone, and the dUTP/dTTP ratio of HAP1 cells. The total [dTTP + dUTP] pool was 61 ± 14 pmol/million cells which was comprised almost entirely of dTTP (55 ± 9 pmol/million cells) and dUTP (7 ± 5 pmol/million cells). Control reactions included polymerase in the absence and presence of added [dUTP + dNTPs] and dUTPase. (**B)** (top) Denaturing urea-PAGE of extension reactions in the presence and absence of dUTPase with cell extract from MDM cells. The image is from one of two biological replicate measurements. (bottom) Total levels of [dTTP + dUTP], dTTP alone, dUTP alone, and dUTP/dTTP ratio of MDM cells. The total [dTTP + dUTP] pool was 0.62 ± 0.04 pmol/million cells which was comprised of nearly equal levels of dTTP (0.32 ± 0.06 pmol/million cells) and dUTP (0.31 ± 0.007 pmol/million cells). Control reactions included polymerase in the absence and presence of added [dUTP + dNTPs] and dUTPase. **(C)** (top) Denaturing urea-PAGE of extension reactions in the presence and absence of dUTPase with cell extract from AM cells. The image is from one of two biological replicate measurements. (bottom) This figure shows the total amount of [dTTP + dUTP], dTTP alone, dUTP alone, and dUTP/dTTP ratio of AM cells. The total [dTTP + dUTP] pool was 0.13 ± 0.005 pmol/million cells, which was comprised of dTTP (0.019 ± 0.004 pmol/million cells) and dUTP (0.11 ± 0.002 pmol/million cells). Control reactions included polymerase in the absence and presence of added [dUTP + dNTPs] and dUTPase.

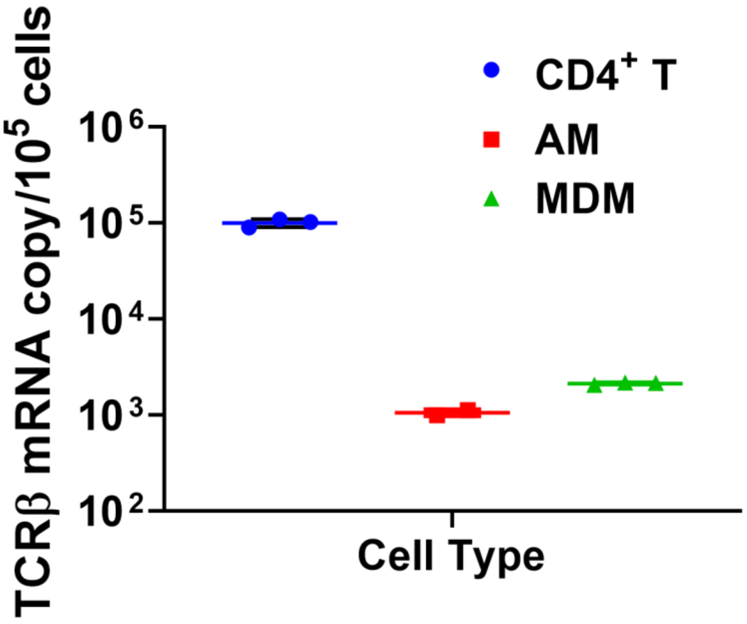


**Figure S2. TCR quantification in various cells.** TCR β mRNA quantification in $CD4^+$ T cells, AM and MDM. *n* = 3 for $CD4^+$ T cells and MDM, *n* = 2 for AM, where *n* is the number of donors.

**Supplemental Table S1. HIV-1 Env V3 amino acid sequences with computational and biological prediction of co-receptor usage**

| Viruses | [a]Env V3 region amino acid sequences | 11/25 rule[b] | Geno2pheno (subtype)[c] | PSSM (scores)[d] | Net Charge[d] |
|---|---|---|---|---|---|
| HXB3 | CTRPNNNTRK**K**IRIQRGPGRAFVT**I**GK-IGNMRQAHC | | | | |
| AM2 | CTRPNNNTRK**S**IHI--GPGRALYT**T**GEIIGDIRQAHC | NSI | R5 (B) | R5 (-11.41) | 5 |
| T2 | CTRPNNNTRK**S**IHI--GPGRALYT**T**GEIIGDIRQAHC | NSI | R5 (B) | R5 (-11.41) | 5 |
| M3 | CTRPNNNTRK**S**IHI--GPGRALYT**T**GEIIGDIRQAHC | NSI | R5 (B) | R5 (-11.41) | 5 |
| T3 | CTRPNNNTRK**S**IHI--GPGRALYT**T**GEIIGDIRQAHC | NSI | R5 (B) | R5 (-11.41) | 5 |
| M4 | CIRPGNNTRR**S**IHI--APGRAFYA**T**GTIIGDIRQAHC | NSI | R5 (B) | R5 (-12.51) | 6 |
| T4 | CTRPSNNTRR**S**IHI--APGRAFYA**T**GAIIGDIRQAHC | NSI | R5 (B) | R5 (-13.07) | 6 |
| M5 | CIRPSNNTRR**S**IHI--APGRAFYA**T**GAIIGDIRQAHC | NSI | R5 (B) | R5 (-13.13) | 6 |
| T5 | CIRPGNNTRR**S**IHI--APGRAFYA**T**GTIIGDIRQAHC | NSI | R5 (B) | R5 (-12.51) | 6 |

[a] Nested PCR products were sequenced using Amplicon-EZ (2 x 250) from each virus, and the Env V3 regions of the most abundant sequence from each sample, aligned to the HXB3 strain, were shown. The positions 11 and 25 of the Env V3 region are indicated in red bold. ‘‘-’’ Represents a sequence gap.

[b] Non-syncytium inducing and syncytium inducing HIV-1 are abbreviated as NSI and SI, respectively.

[c] CCR5- and CXCR4- tropic viruses are designated R5 and X4, respectively.

[d] Calculations done using https://indra.mullins.microbiol.washington.edu/webpssm

**Supplemental Table S2. Hypermutation analysis of the most abundant sequence from each QVOA[a]**

| Sequence | Muts: (Match Sites) | Out of: (Potential Mut Sites) | Controls: (Control Muts) | Out of: (Potential Controls) | Rate Ratio: (A/B)/(C/D) | P-value: = P (Muts, Poten. Muts - Muts, Cntrls, Poten. Cntrls - Cntrls)[b] |
|---|---|---|---|---|---|---|
| **A2(49.77%)** | 0 | 25 | 3 | 19 | 0.00 | 1 |
| **T2(14.01%)** | 0 | 25 | 3 | 19 | 0.00 | 1 |
| **M3(61.33%)** | 0 | 25 | 3 | 19 | 0.00 | 1 |
| **T3(23.06%)** | 0 | 25 | 3 | 19 | 0.00 | 1 |
| **M4(20.08%)** | 2 | 25 | 1 | 19 | 1.52 | 0.604047 |
| **T4(40.13%)** | 0 | 25 | 1 | 19 | 0.00 | 1 |
| **M5(66.25%)** | 2 | 25 | 0 | 19 | inf | 0.317125 |
| **T5(5.44%)** | 2 | 25 | 0 | 19 | inf | 0.317125 |

[a]The pattern definitions are as follows, where No pattern is indicated as '…':

| Pattern | Upstream | From | →To | Downstream |
|---|---|---|---|---|
| 'Mut' | … | G | →A | RD … |
| 'Control' | … | G | →A | YN\|RC … |

[b]'Potential Mut' or 'Potential Control' means a match to the corresponding Upstream, From, and Downstream patterns above, while an actual 'Mut' matches those and the To pattern as well. Using the default settings, a P-value less than 0.05 is the cut-off to indicate a hypermutation site.

### Supplemental Table S3. Primers, probes and oligos used in this study

| Gene | Forward sequence (5′–3′) | Reverse sequence (5′–3′) |
|---|---|---|
| **UNG2** | GCCAGAAGACGCTCTACTCC | TCGCTTCCTGGCGGG |
| **APE1** | TGGAATGTGGATGGGCTTCGAGCC | AAGGAGCTGACCAGTATTGATGA |
| **Polβ** | GGCAGTTTCAGAGGTGC | GGCAAACACCCATGAACTTT |
| **LIG III** | GATCACGTGCCACCTACCTTGT | GGCATAGTCCACACAGAACCGT |
| **DUT** | GGGAGAATCACTTGAGGTTGAG | GGGTTCTCTCTCTCCTTCTCTT |
| **SAMHD-1** | GGATTACTAAAAACCAGGTTTCACAACT | TGTCGTTCCATTCCTTTTTTTGA |
| **18s rRNA** | TGTGCCGCTAGAGGTGAAATT | TGGCAAATGCTTTCGCTTT |
| **ERT** | GCTAACTAGGGAACCCACTGCTT | CAACAGACGGGCACACACTGCTT |
| **ERT probe** | FAM-AGCCTCAATAAAGCTTGCCTTGAGTGCTTC-BHQ2 | |
| **LRT** | TGTGTGCCCGTCTGTTGTGT | GAGTCCTGCGTCGAGAGATC |
| **LRT probe** | FAM-CAGTGGCGCCCGAACAGGGA-BHQ2 | |
| **Alu** | GCCTCCCAAAGTGCTGGGATTACAG | |
| **Gag** | CATGTTTTCAGCATTATCAGAAGGA | TGCTTGATGTCCCCCCACT |
| **Gag probe** | FAM-CCACCCCACAAGATTTAAACACCATGCTAA-BHQ2 | |
| **RPP30** | GATTTGGACCTGCGAGCG | GCGGCTGTCTCCACAAGT |
| **RPP30 probe** | VIC-CTGACCTGAAGGCTCT-MGBNFQ | |
| **TCR β (V)** | ACACGTGAAATGCTCTTTGCG | TTACTCCTGCGCCTCTGTGTC |
| **TCR β (C)** | TGGCTTCTGGCACTCCTTG | GCCATGTGAAGACAGAGGCA |
| **E90 (Out-FWD)** | CACAGTACAATGTACACATGGAAT | |
| **Nesty8 (Out-REV)** | | CATACATTGCTTTTCCTACT |
| **DLoop (In-FWD)** | GTCTAGCAGAAGAAGAGG | |
| **E115 (In-REV)** | | AGAAAAATTCCCCTCCACAATTAA |
| **SNE probe** | 5′FAM-TGTTCTATGTTCATACACCACA-3’ | |
| **SNE template** | 3′-ACAAGATACAAGTATGTGGTGT**A**-5’ | |

**Supplemental Table S4. Characterization of targeted amplicon sequencing data**

| Sample ID[a] | | Barcode Sequence | # Reads | Mean Quality Score[b] | % Bases >= 30 | Major Variant (%)[b] |
|---|---|---|---|---|---|---|
| **P2** | AM2 | GCGTAGTA+AGGCTATA | 533,214 | 35.46 | 91.11 | 49.77 |
| | T2 | GCGTAGTA+GCCTCTAT | 349,159 | 35.42 | 90.89 | 14.01 |
| **P3** | M3 | GCGTAGTA+AGGATAGG | 324,326 | 35.5 | 91.27 | 61.33 |
| | T3 | AAGCGACT+TCAGAGCC | 460,898 | 34.98 | 88.44 | 23.06 |
| **P4** | M4 | AAGCGACT+TAAGATTA | 335,959 | 35.04 | 88.74 | 20.08 |
| | T4 | AAGCGACT+CTTCGCCT | 258,513 | 35.08 | 88.9 | 40.13 |
| **P5** | M5 | AAGCGACT+GTCAGTAC | 390,088 | 34.96 | 88.34 | 66.25 |
| | T5 | AAGCGACT+ACGTCCTG | 381,607 | 34.94 | 88.24 | 5.44 |

[a]P refers to patient, AM is alveolar macrophage, M is monocyte, T is $CD4^+$ T cells

[b]The percentage calculated as the (# of reads of the variant/total reads) X 100
